# Cognitive exertion affects the appraisal of one’s own and other people’s pain

**DOI:** 10.1101/2022.06.09.495450

**Authors:** Laura Riontino, Raphael Fournier, Alexandra Lapteva, Nicolas Silvestrini, Sophie Schwartz, Corrado Corradi-Dell’Acqua

## Abstract

Correctly evaluating others’ pain is a crucial prosocial ability, especially relevant for the healthcare system. In clinical settings, caregivers assess their patients’ pain under high workload and fatigue, often while dealing with competing information/tasks. However, the effect played by such cognitive strain in the appraisal of others’ pain remains unclear. Following embodied accounts that posit a shared representational code between self and others’ states, it could be hypothesized that the representation of people’s pain might be influenced by cognitive exertion similarly to first-hand experiences.

Fifty participants underwent one of two demanding tasks, involving either working memory (Experiment 1: N-Back task) or cognitive interference (Experiment 2: Stroop task). After each task, participants were exposed to painful laser stimulations at three intensity levels (low, medium, high), or video-clips of patients experiencing three intensity levels of pain (low, medium, high). Participants rated the intensity of each pain event on a visual analogue scale.

We found that the two tasks influenced rating of both one’s own and others’ pain, by decreasing the sensitivity to medium and high events. This was observed either when comparing the demanding condition to a control (Stroop), or when modelling linearly the difficulty/performance of each depleting task (N-Back).

We provide converging evidence that cognitive exertion affects the subsequent appraisal of one’s own and likewise others’ pain. Healthcare personnel should be aware that high workload might alter their cognitive abilities.

**Perspective:** this research shows that cognitive effort aftereffects impact negatively the assessment of of medium/high pain in others, reminiscently to what was observed in first-hand experiences. Healthcare professionals should be aware that high workload and severe cognitive fatigue could affect their diagnostic skills.

## Introduction

Identifying and assessing the intensity of pain experienced by others is a crucial ability, in both clinical and private environments^3,37,57–59^. Unfortunately, caregivers, like physicians/nurses or individuals in full-time charge of suffering family members, might be subjected to fatigue, poor sleep and long/irregular work shifts that may overload of their cognitive resources. However, it is still unclear how extensive mental exhaustion might interfere with the assessment of others’ pain. One plausible effect could be linked to the known tendency of caregivers to underestimate^57,58^ (and consequently undertreat^59^) the pain of people under their care, a phenomenon that becomes more pronounced with longer experience in the field ^16–18,24,26^. Unfortunately, to our knowledge, information about the role played by cognitive exertion in the lay population (unbiased by long-terms exposure in clinical settings) is lacking. In this study we sought to investigate pain assessment under the lens of the well-known “ego depletion” paradigm, which is linked to a lack of cognitive resources associated with decreased performance due to a previous exertion of self-control^32,85^.

Previous studies converge in finding that depletion of individual cognitive resources led to a generally reduced sensitivity to *concurrent* own pain experience^11,50,70,79^. However, it is less clear whether similar influence linger also *after* the task completion. For instance, being engaged in a highly demanding Stroop task decreased the sensitivity to subsequent intense stimulations in a study^33^, but it increased the sensitivity to mild noxious events in others^65,68^. Such mixed findings beg for a systematic investigation of after-effects of cognitive exertion on different pain levels, and across different depleting tasks, to account also for concerns about reliability of such paradigm which have recently been put forward by the research community^29,41^.

Most critically, it is unclear whether cognitive exertion would impact the assessment of pain of others similarly to one’s own. Seminal models from social psychology/neuroscience suggest that others’ suffering is partially processed in an *embodied* fashion, i.e. by recruiting the same mechanisms underlying one’s own first-hand experiences^1,5,15,30,31^. Indeed, analgesic manipulations (e.g., acetaminophen, hypnosis, placebo) can similarly influence the sensitivity to self and others’ pain^8,49,60^. Furthermore, neuroimaging studies suggest that self and others’ pain share a partly-common neuronal representation in a widespread network including the middle cingulate cortex^8,21,22^, a brain region held to play a key role in the regulation of one’s pain responses^63,65^ (but see^38^). Hence, it is reasonable to assume that such regulatory mechanisms might equally impact the representation of one’s and others’ pain.

An alternative model, the *broaden-and-build* theory, suggests instead that positive emotions/mood can broaden one’s resources, improving physical, intellectual and social abilities, whereas negative emotions would narrow one’s resources, thus promoting self-related thoughts^28^. Consistently, previous studies showed that negative events cause hyperalgesia^4,7,25,42,48,54,56,76,81–83,86^, but have opposite effects for the pain of others by decreasing physiological and neural response to the sight of people’s injuries^54^. Importantly, the *broaden-and-build* theory explains these effects in terms of an indirect modulation of emotions on one’s personal resources^28^. Consequently, narrowing one’s resources through other means (e.g., cognitive exertion) should lead to the same effects, by increasing the sensitivity to self-pain (as found by^65,68^) at the expense of sensitivity to other people’s pain.

Here we engaged participants in one of two demanding paradigms, involving either working memory (Experiment 1: N-Back task) or inhibition (Experiment 2: Stroop task), to investigate the effects of cognitive exertion on the subsequent assessment of one’s and others’ pain. Based on previous research, we expected the demanding tasks to influence intensity ratings of self-pain, albeit with unascertained direction^26,62,65^. The critical question was whether the same manipulation would influence the pain perceived in others in similar (as predicted by *embodied* accounts) or opposite fashion (suggested by the *broaden-and-build* theory) with respect to self-experienced pain.

## Experiment 1

### Methods

#### Participants

Recruitment took place through advertisements posted at the University of Geneva buildings and online platforms. We excluded from our experiment all individuals with the following characteristics: history of substance/alcohol abuse, history of neurological or psychiatric illness, history of chronic pain, fever or an ongoing acute medical condition and extreme dark skin colour (because radiation absorption of dark skin is higher for the wavelength of the laser device used for nociceptive stimulations^2,46^). They declared good health and typical cognitive proficiency, they had good (or corrected) visual acuity. Based on these criteria, we recruited a total of 36 participants, ten of which were subsequently excluded due to lack of sensitivity to the nociceptive stimulation). The latter criterion was established in those participants who, during the main experiment, rated the highest stimulation under the control condition < 4 on a 10-points pain intensity scale (see below for more details), despite good sensitivity to pain during a preliminary calibration phase. We further excluded 1 further participant who was not susceptible to the task manipulation. This was achieved by calculating a combined index for accuracy and response time (Inverse Efficiency Score^55,72^, see data processing subsection) during the task, and by removing the subject whose performance in the difficult (demanding) condition was comparable/better than in the control condition (see Results section and Figure 1, left subplot). This led to a final sample of N = 25 (14 females; aged 19 to 37, Mean = 25.16, SD = 5.03 years). Sample size was consistent with the previous studies on which the present research is based^12,68^. Participants were all naïve to the purpose of the study and none of them participated in both experiments. This study was approved by the local ethical committee (*Commission Cantonale d’Éthique de la Recherche* of Geneva, protocol code: CCER N. 2019-01355) and conducted according to the Declaration of Helsinki.

**Figure 1.**
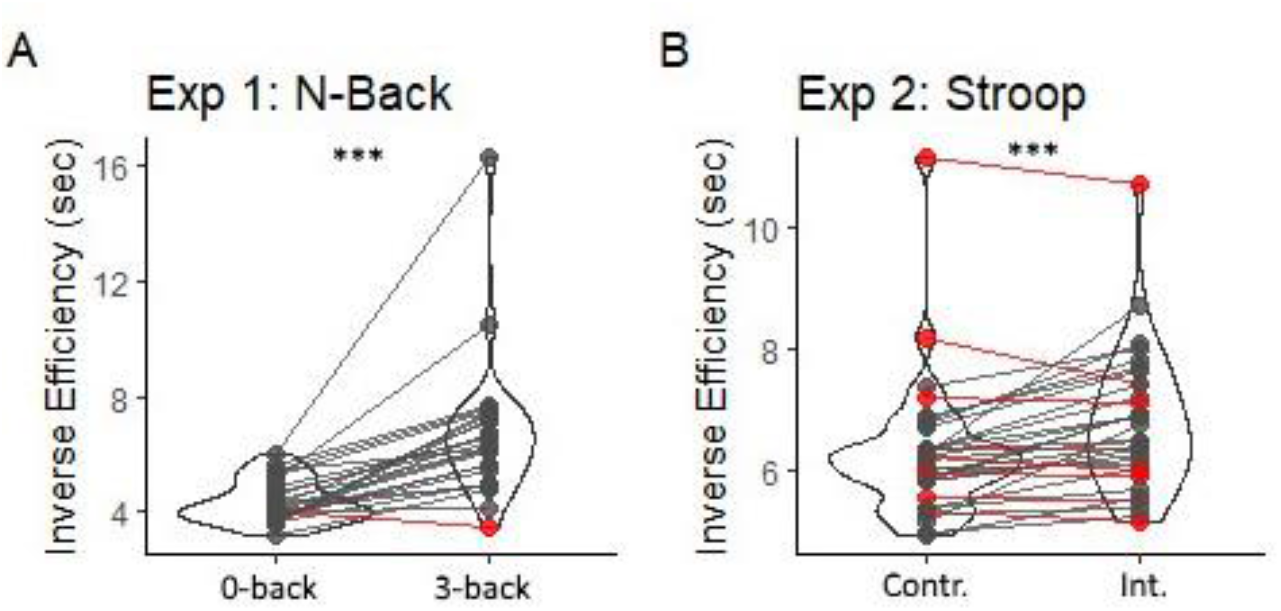
Violin plots and individual inverse efficiency data describing the difference between the two main conditions in **(A)** the N-Back (Exp. 1) and **(B)** Stroop task (Exp. 2). *** p < 0.001 for the effect of task difficulty. Inverse Efficiency scores refer to Response Times of correct responses (sec) penalized proportionally to the number of errors performed in the same condition. Red dots/lines refer to those specific participants for whom the difficult condition was associated with comparable/lower scores than the control condition. Int. = Stroop Interference. Contr. = Stroop Control.

#### Experimental Set-up

To disentangle the opposing theoretical predictions concerning the impact of exertion on others’ pain judgment, volunteers assessed their pain sensitivity and judgments of others’ pain after a low or high cognitive demanding N-Back task. Within this framework, we systematically modulated workload (factor Load: low vs high), type of pain (factor Type: self vs other) and pain intensity (factor Intensity: low, medium, high).

##### N-Back task

We administered an N-Back task^35^, which was previously used successfully to evaluate the impact of distraction on pain sensitivity^12^. Participants monitored a series of stimuli appearing one at a time on the screen. They were required to identify if the stimulus presented was the same as the one presented *n* previous items by pressing one button (‘yes’) and if not by pressing a different button (‘no’). In the current study, *n* was set to be 3 (high cognitive load condition) or 0 (low cognitive load condition). In the 3-Back condition, participants were required to constantly update the target letter, by retrieving from working memory the one letter occurring 3 items before. For instance, if we pretend a sequence of letters “RGSR”, correct answer to the last would be “yes”, as the same letter occurred also three trials before in the sequence “RSGP” but in this case answer is “no”. As control, we implemented a low cognitive load condition, in which participants were asked to respond only on the trial that they were currently processing, by pressing “yes” whenever a predefined stimulus (“V” letter) would be presented. In each block, a total of 18 letter stimuli were presented one at a time with 4 target-hits.

Each letter was presented for 500 ms, followed by an inter-trial interval (ITI). The ITI between letters of the 0-Back condition was fixed at 700 ms. Instead, for the 3-Back condition to reduce within-participant variation, the ITI was adapted based on participants’ performance in the previous block (i.e., corresponding to 18 previous trials). This is roughly comparable to a previous study with a similar design (adaptation based on previous 15 trials^12^), although in that case, the authors tested pain concurrently to the N-Back, and therefore had no need to divide the experimental session into small blocks^12^. Target sensitivity was assessed with the non-parametric signal detection measure A’ ^69,71^. This parameter is computed from a pair of hit and false alarm rates^12^: a value near 1 indicates a good discriminability, whereas a value near 0.5 (when hits are equal to false alarm) indicates chance performance. Initial duration of the empty screen was 700 ms. Then based on the previous trial, when discriminability was higher than the targeted level of A’ = .75, empty screen interval was reduced, while with a discriminability lower than A’ = .75 it was increased.

Prior to the main experiment, participants carried out a brief training session, in which the order of the blocks was fixed: the first block was 0-Back, the second block was 3-Back in which subsequent adjustments were made after each trial. Instead, in the main experiment, the order of blocks was pseudorandomized, and difficulty of the task was calibrated by adapting the interval for each participants’ performance to the previous 3-Back block. Thirteen participants (7 women) started with two blocks 0-Back condition (low cognitive load) and then two blocks with high cognitive load (combined with 2 “Self” and 2 “Other” in a pseudorandomized order) followed by a break of about 5 minutes and then the same task sequence was repeated (combined with 2 “Self” and 2 “Other” in a pseudorandomized order). With the same procedure, twelve participants (7 women) started with the 3-Back condition (high cognitive load).

##### Self-Pain

Half of the N-Back blocks were followed with painful stimulations on their own body. Self-pain was administered through nociceptive radiant heat stimuli was delivered by an infrared neodymium: yttrium-aluminum-perovskite laser system (Neodinium: Yttrium Aluminium Perovskite; Stimul 1340 El.En®; wavelength 1.34 µm; pulse duration 4 ms, beam diameter 6 mm^2^). At this short wavelength, the skin is highly transparent to laser stimulation, and consequently, the laser pulses directly and selectively activate Aδ and C fibers nociceptive terminals located in the superficial layers of the skin^34,53^. Since these fibers have different conduction velocity, participants experienced a double sensation: an initial pricking pain due to Aδ-fiber stimulation, followed by a C-fiber-related burning pain. Laser pulse was transmitted through an optic fiber of 10-meter length, with a diameter of 6 mm by focusing lenses. Laser beam was directed to a rectangular skin area (approximately 4 × 2 cm, main axis mediolateral) on the back of the non-dominant hand. To avoid receptor sensitization, as well as damages from long-term exposure, the laser beam was relocated after each trial within a predefined stimulated skin area (see also^43^).

To ensure protection from adverse effects of the laser, the following procedure were put in place. The experiment was carried out in an *ad hoc* room, protected with laser safety curtains certified for the wavelength of the stimulator. Furthermore, participants and experimenters wore eye-protection goggles with optical density ≥ 2 at 1.34 µm. Participants were asked to remove all accessories from their non-dominant hand to allow safe stimulation of the skin area.

Prior to the beginning of the experiment, pain stimulus intensity was individually calibrated for each participant based on a random staircase thresholding session^45^ to determine stimulations eliciting three levels of pain that were then used in the main experiment. Each trial started with a flash appearing on the screen for 3 seconds indicating that the stimulation was about to come, while the laser was preparing to deliver the correct amount of energy. After each stimulation, participants answered on a 10-point visual analogue scale ranging from not at all painful (0) to worst pain imaginable (10). Extreme manikins of the arousal scale of the Self-Assessment Manikin (SAM) were displayed at the extremities of the scale for assessing pain intensity as an economical and straightforward cue^47^. We also included numbers under the rating line, coloured from green (0) to red (10), to visually help participants choosing their answer. In this context, participants were instructed to consider the landmark “4” on the scale as the ideal point where they started to feel the stimulation as painful. The rating was self-paced with no time restrictions. The calibration session lasted about 15 minutes. At the end of this session, we obtained a pain curve based on which we would then select three different energies that corresponded to three different levels of pain for each participant for low (L), medium (M) and high (H) intensity respectively (medians of about 2, 4 and 6 on the pain rating scale).

In the main experiment, after each corresponding N-Back block, participants were exposed to six stimulation-trials in random order. These were characterized by 2 Low, 2 Medium and 2 High pain stimulations. Each trial started with a flash appearing on the screen for 3 seconds indicating that a stimulation was about to come. The nociceptive stimulus was followed by a black screen lasting for 1 second and then the visual analogue scale for pain judgement. This was identical to the one used in the calibration session, except for the absence of intermediate tick lines, and a time constraint of 5 seconds for providing a response.

After the pain intensity rating, a 1 second of black screen was presented, followed by a second rating scale. This was a pain metacognition scale probing for participants’ confidence about their previous rating. In particular, participants had to express their confidence about their previous pain judgment on a 10-point visual analogue scale ranging from no confidence at all (0) to completely confident (10). Extreme manikins of the dominance scale of the Self-Assessment Manikin (SAM) were included at the extremities of the metacognition scale^47^. As for the case of pain intensity ratings, participants had 5 seconds also to provide a rating. This second scale was introduced to assess a secondary hypothesis that cognitive exertion could influence, not only participants’ pain assessment, but also the degree of which they were aware of such potential influence.

Following the confidence assessment, a black screen appeared for a random inter-trial interval ranging between 2.5 and 4.5 seconds with steps of 200 ms. Overall, each block, and subsequent pain stimulations, lasted about 3 minutes.

##### Other-Pain

The remaining half of the blocks were followed by saw video-clips describing pain in others. For this purpose, we used stimuli taken from a database of videos of patients faces, which had spontaneous (non-simulated) expressions of pain^44^. Thirty videos were cut to last 3 seconds and to show the most salient expressions. For video piloting, 21 independent participants rated these short videos online (14 women; age 22 to 35, Mean = 27.6, SD = 4.7). For the purpose of this experiment, we selected 24 videos based on median ratings of the piloting that could match the three levels of self-pain (8 videos per each level of pain), one characterized by low (3 women, 5 men; Mean = 0.58, SD = 0.50), one characterized by medium (4 women, 4 men; Mean = 3.12, SD = 0.60) and one characterized by high (4 women, 4 men; Mean = 4.70, SD = 0.49) level of painful expression. Furthermore, there was high agreement between the 21 raters in their assessment of the videos of each pain level, as shown by the average inter-rater-correlation (low pain, Spearman’s *ρ* = 0.40 [95% CI: 0.15, 0.67]; medium, *ρ* = 0.50 [0.11, 0.70]; high, *ρ* = 0.65 [0.40, 0.82]) and Cronbach’s α (low, α = 0.81 [0.61, 0.89]; medium, α = 0.91 [0.79, 0.96]; high, α = 0.93 [0.85, 0.96]). Finally, as the people depicted in the videos were also probed to rate their own pain experience^44^, we could thus check that the videos for the three levels of other’s pain exhibited clear-cut differences in low, medium and high self-reported pain. This insured that the selected clips clearly depicted three distinct intensity levels from both the point of view of the video-recorded people, and from an independent sample of observers.

##### Procedure

The experiment was conducted under the following procedure. On arrival, participants read carefully and signed the informed written consent, then they were reminded that each laser stimulus would be very fast (only 4 ms) and that they could ask to stop the experiment at any time. All tasks and stimuli were coded, managed and presented with Psychophysics Toolbox Version 3 (PTB-3, http://psychtoolbox.org/), a free set of Matlab R2018b (Mathworks, Natick, MA). The experimental session consisted of two parts: 1) pre-test divided in pain thresholding session and training of the N-Back task, and 2) main experiment. Participants had to use a response box with four buttons in their dominant hand to answer. Two experimenters were always present during the experimental session. One experimenter was responsible for technical checks of the equipment before and during acquisition, including physiological measures, laser inputs and behavioural recordings. The other experimenter was responsible for interacting with participants and delivering painful stimulations by directing the laser beam on participants’ hand. Importantly, the latter experimenter was unaware of the level of pain emitted by the laser.

In the main experiment, each participant performed four blocks representing the 3-Back (cognitive exertion) and four blocks of a 0-Back (baseline), presented in a pseudorandom order. Each block was followed the delivery of the six pain stimulation stimulation-trials delivered to either the self (through nociceptive laser stimulations) or the other (through video-clips). This led to a 2 (Task: 3-Back *vs*. 0-Back) x 2 (Type: Self *vs*. Other) structure of each block, within each which are administered pain events of 3 intensity levels (low, medium and High). Overall, the experiment session lasted 1 hour and half and, at the end, participants were refunded for their time and effort.

##### Questionnaires’ battery

Following the main experimental session, participants were asked to fill some questionnaires. In addition to the general information sheet (including questions about sex, age and hand dominance), participants filled out the Pain Catastrophizing Scale^73^ (PCS) and the Interpersonal Reactivity Index^23^ (IRI), because we hypothesized that these dimensions of personality might modulate the effects of interests. In the PCS, participants were asked to indicate the degree to which they think and feel when they experience pain using a 0 (not at all) to 4 (all the time) scale. A total score is yielded (ranging from 0-52), along with three subscale scores assessing rumination, helplessness and magnification, which respectively report rumination about pain (e.g. “I can’t stop thinking about how much it hurts”), feeling helpless to manage pain (e.g. “There is nothing I can do to reduce the intensity of my pain”) and magnification of pain (e.g. “I’m afraid that something serious might happen”).

The IRI measures various aspects of empathy including cognitive and emotional empathy. Participants had to answer 28 statements using a 5-point scale, ranging from “Does not describe me well” to “Describes me very well”. This measure is divided in four seven-item subscales. The perspective taking (PT) scale reports tendency to spontaneously adopt the psychological point of view of others in daily life (e.g. “I sometimes try to understand my friends better by imagining how things look from their perspective”). The empathic concern (EC) scale assesses the tendency to experience feelings of sympathy and compassion for unfortunate others (e.g. “I often have tender, concerned feelings for people less fortunate than me”). The personal distress (PD) scale taps the tendency to experience distress and discomfort in response to extreme distress in others (e.g. “Being in a tense emotional situation scares me”). The fantasy (FS) scale measures the tendency to imaginatively transpose oneself into fictional situations (e.g. “When I am reading an interesting story or novel, I imagine how I would feel if the events in the story were happening to me”).

#### Data processing

##### Behavioral Measures

Despite all 36 participants exhibited good sensitivity to pain during the calibration phase, effects of habituation/desensitization might occur in the main experiment. For this reason, we inspected the average rating of the most intense laser stimulation following those blocks which were not held to induce cognitive exhaustion (i.e., 0-Back). We did not include the difficult condition (3-back) as in those blocks sensitivity could have been dampened due to the main manipulation. For ten participants the rating was < 4 on the 10 points intensity scale. As we instructed to consider 4 as the ideal point where they started to feel the stimulation as painful (see above), those participants were excluded.

We then checked whether participants were susceptible to the N-Back task. To this purpose, we considered all participants who displayed good sensitivity to pain (N = 26). In this analysis, trials with reaction times < 100 ms were removed as potentially reflective anticipatory response unrelated to the task. For each participant, each task and each condition, we calculated the average accuracy, median reaction times of correct responses, and combined Inverse Efficiency Score (IES), with 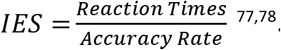. This combined measure refers to Reaction Times of correct responses *penalized* proportionally to the number of errors performed in the same condition (e.g., if a given condition is associated with an accuracy of 90%, IES corresponds to correct Reaction Times multiplied by ∼10%), and aims at maximizing the information from those cases in which correct Reaction Times and Accuracy are negatively coupled (i.e., conditions eliciting the slowest correct responses being also the least accurate). Omitted responses were considered as errors. After acquisition, all these measures were fed to a paired-sample t-test assessing effects of task difficulty (3-Back *vs*. 0-Back). Then, individual data were used to exclude from all subsequent analyses those participants who were not susceptible to the task (final sample N = 25 see Results and Figure 1, left subplot).

Having established participants’ susceptibility to the task, we tested its effect on the subsequent pain intensity ratings. This was achieved though a linear mixed model with task condition (3-Back vs 0-Back) and intensity of pain (low, medium, high) as fixed factors. In this case, omitted responses were removed from the analysis. The subject’s identity was modelled as random factor, with random intercept and slope for the two fixed factors and the interaction therefor. Specifically, we ran two models, one for the Self-Pain condition, and the second for the Other-pain conditions. Significance associated with the fixed effects was calculated with a Type III Analysis of Variance using the Satterthwaite approximation of the degrees of freedom, as implemented in the *lmerTest* package^39^ of R (https://cran.r-project.org/) software.

##### Physiologic Responses

Since it has been shown that cognitive exertion has a negative carryover effects on physical performance^9^ and that physiological states correlate with cognitive fatigue^40^, physiological measures during pain stimulation were also collected to examine possible subsequent effects. Therefore, during the main experiment, we continuously recorded physiological responses to stimuli from their left hand, such as skin conductance (EDA) and heart rate. Detailed information about the analysis of these measures in provided in Supplements.

## Results

### Task Demands

We first assessed whether participants exhibited higher difficulty in the demanding conditions. We focused on all participants who displayed adequate sensitivity to pain (N = 26; see Methods) and found that, on average, they displayed significant differences across the two conditions of interest in accuracy (*t*_(25)_ = -8.64, *p* < 0.001; *d* = -1.73), reaction times of correct responses (*t*_(25)_ = 5.45, *p* < 0.001; *d* = 1.09), and combined inverse efficiency score (IES, *t*_(25)_ = 5.83, *p* < 0.001; *d* = 1.17). However, although the tasks appeared to influence the performance of the overall population, one participant appeared non-susceptible to the manipulation, as one participant showed more proficient performance in the difficult condition (Figure 1, left subplot). Because the goal of the study was to test for the effect of increased task demand on pain processing, the data from this participant was excluded from subsequent analysis, because it did not establish higher demand for the more difficult versions of the tasks. We thus obtained a final sample of N = 25.

### Pain Ratings

We then assessed whether sensitivity to nociceptive stimulation on one’s own body was influenced by the preceding task. Table 1 reports results associated with pain intensity ratings, and revealed a significant main effect of stimulation *Intensity*, confirming that participants rated the three intensity levels as progressively more painful. No modulation of *Task* was observed, neither as a main effect, nor in interaction with the *Intensity*. We also checked whether participants’ sensitivity to one’s pain changed as function of the performance of the preceding 3-Back condition. This was achieved by modelling participants’ inverse efficiency score (IES, see Methods) from the previous block as a between-subjects continuous predictor. We chose this measure as it represents an efficient combination of correct Response Times and Accuracy values, which prevents us from running two separate analyses. Table 2 and Figure 2A reports the results of the main analysis, confirming the main effect of stimulation *Intensity*, but also describing a significant *IES*Intensity* interaction. In particular, the interaction reveals that blocks associated with poorer performance led to significant decreased sensitivity to MP (*vs*. LP) stimulations. A similar interaction was observed also at the level of HP (*vs*. LP), although only at a marginal level (Table 2). Finally, as the 3-Back task was adjusted online to stabilize performance across blocks (see Methods), IES describes only part of the inter-individual variability in performance, whereas the remaining part of the information is provided by the difficulty parameter (the inter-trial interval, ITI, see methods) of the task and how they changed from block to block. We therefore also modelled responses as function of the median ITI from the preceding block. However, such parameter did not affect participants’ sensitivity to pain, neither as main effect nor in interaction with intensity (see Table 2).

**Table 1.**
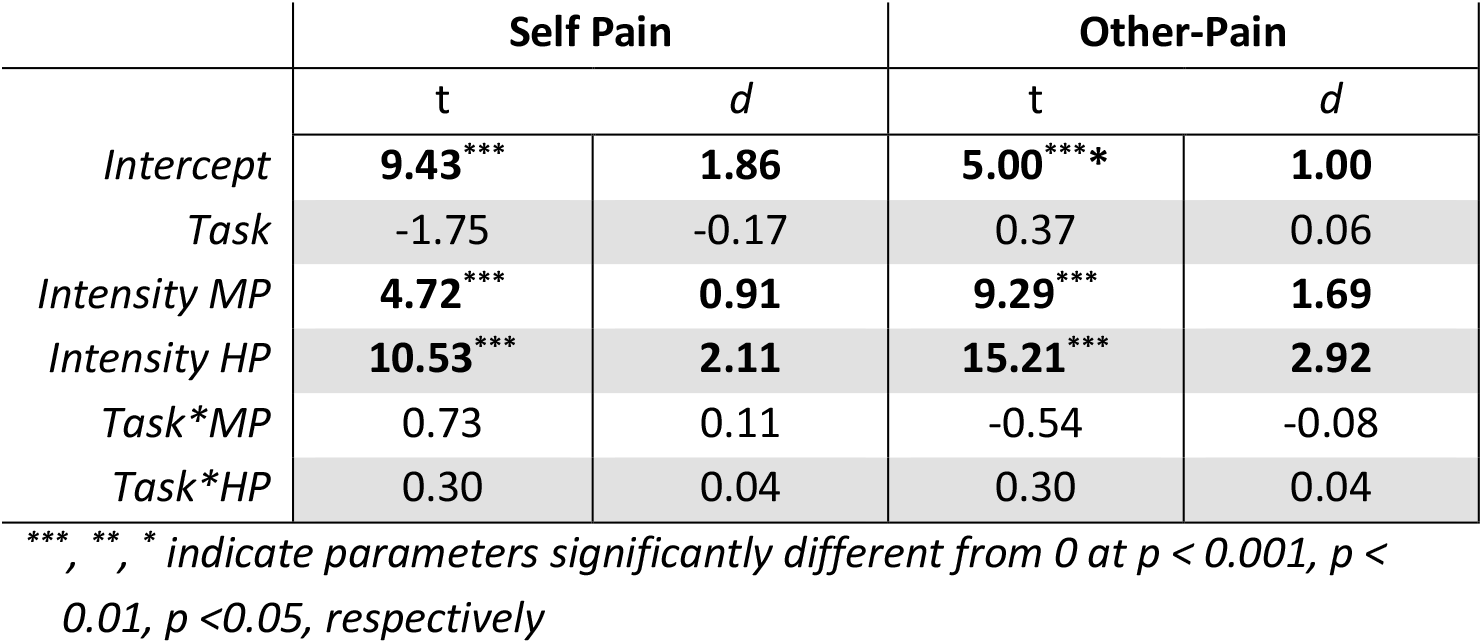
Experiment 1. Analysis of Task-effects on pain. We report the t-values and effect sizes (d) associated with parameter estimates from linear mixed model analyses testing effects of task on Self and Others’ pain. Significant effects are highlighted based on the corresponding p-value. MP = Medium Pain; HP = High Pain.

|  | Self Pain |  | Other-Pain |  |
| --- | --- | --- | --- | --- |
|  | t | d | t | d |
| *Intercept* | **9.43<sup>\*\*\*</sup>** | **1.86** | **5.00<sup>\*\*\*</sup>** | **1.00** |
| *Task* | -1.75 | -0.17 | 0.37 | 0.06 |
| *Intensity MP* | **4.72<sup>\*\*\*</sup>** | **0.91** | **9.29<sup>\*\*\*</sup>** | **1.69** |
| *Intensity HP* | **10.53<sup>\*\*\*</sup>** | **2.11** | **15.21<sup>\*\*\*</sup>** | **2.92** |
| *Task\*MP* | 0.73 | 0.11 | -0.54 | -0.08 |
| *Task\*HP* | 0.30 | 0.04 | 0.30 | 0.04 |

<sup>\*\*\*</sup>, <sup>\*\*</sup>, <sup>\*</sup> indicate parameters significantly different from 0 at $p < 0.001$ , $p < 0.01$ , $p < 0.05$ , respectively

**Table 2.** Experiment 1. Analysis of Task Performance. We report the t-values and effect sizes (d) associated with parameter estimates from linear mixed model analyses testing effects of task performance and difficulty on Self and Others’ pain. Significant effects are highlighted based on the corresponding p-value. MP = Medium Pain; HP = High Pain; IES = Inverse Efficiency Score; ITI = Inter-trial Interval; Cov. = Covariate of Interest (IES, ITI).

|  | Self-Pain: IES |  | Other-Pain: IES |  | Other-Pain: ITI |  |
| --- | --- | --- | --- | --- | --- | --- |
|  | *t* | *d* | *t* | *d* | *t* | *d* |
| *Intercept* | **9.33<sup>\*\*\*</sup>** | **1.85** | **6.11<sup>\*\*\*</sup>** | **1.22** | **6.22<sup>\*\*\*</sup>** | **1.21** |
| *Intensity MP* | **6.17<sup>\*\*\*</sup>** | **1.07** | **9.18<sup>\*\*\*</sup>** | **1.57** | **9.47<sup>\*\*\*</sup>** | **1.43** |
| *Intensity HP* | **10.81<sup>\*\*\*</sup>** | **2.23** | **13.43<sup>\*\*\*</sup>** | **2.81** | **14.02<sup>\*\*\*</sup>** | **2.84** |
| *Covariate* | 0.94 | 0.09 | 1.90 | 0.19 | 1.50 | 0.10 |
| *Cov. \*MP* | **-2.81<sup>\*\*</sup>** | **-0.20** | **-2.89<sup>\*\*</sup>** | **-0.24** | **-2.50<sup>\*</sup>** | **-0.17** |
| *Cov. \*HP* | -1.76<sup>^</sup> | -0.15 | -0.95 | -0.10 | -1.73 | -0.12 |

<sup>\*\*\*</sup>, <sup>\*\*</sup>, <sup>\*</sup> indicate parameters significantly different from 0 at $p < 0.001$ , $p < 0.01$ , $p < 0.05$ , respectively.
<sup>^</sup> indicate effect at the marginal ( $0.05 < p < 0.1$ ).

**Figure 2.**
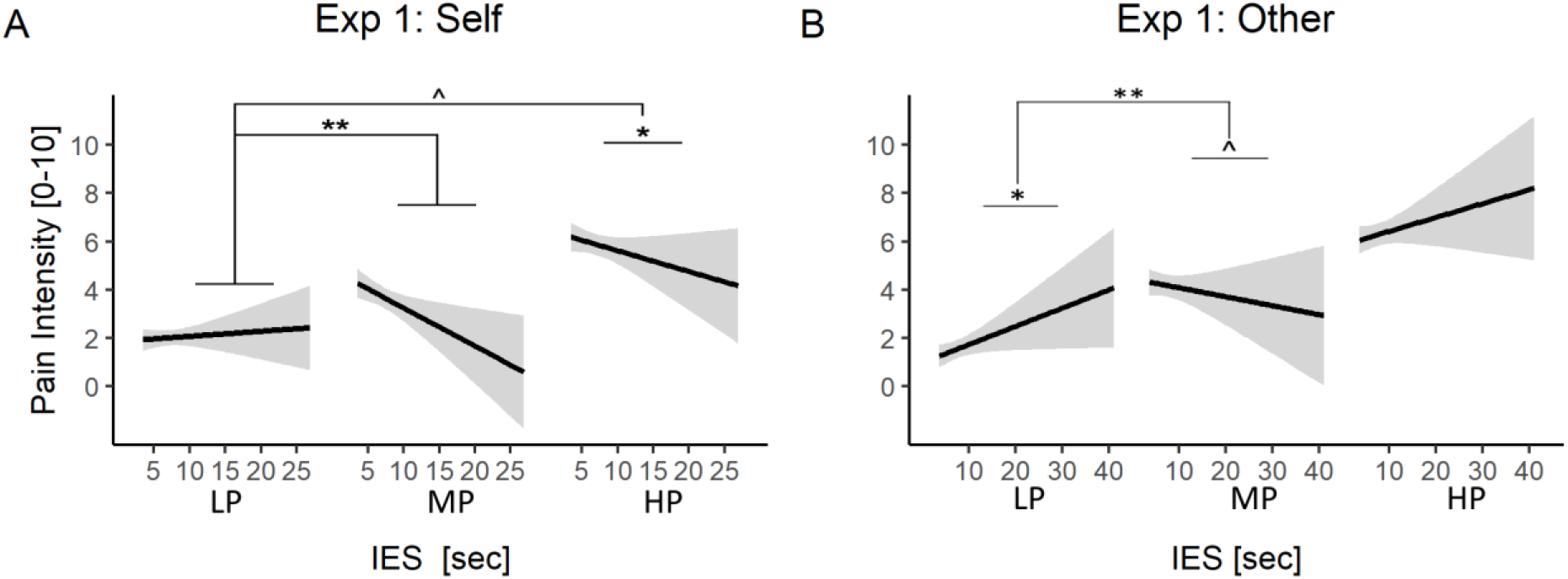
Experiment 1. Analysis of Task Performance. Performance effects on Pain Intensity Stimulation results associated with the N-Back task for Self-Pain **(A)** and Other-Pain data **(B)**. For each pain level, the relationship between IES and pain ratings is described through a linear regression line with 95% confidence interval area. “**”, “*”, “^” refer to simple IES effects for a given pain level, or significant interactions between IES and pain intensity at p < 0.01, p<0.05, and 0.05<p<0.1, respectively. IES = Inverse Efficiency Score; LP = Low Pain; MP = Medium Pain; HP = High Pain.

Next, we analysed pain ratings associated with videos of patients displaying painful/painless facial expressions. Results are reported in Table 1 and reveal a significant main effect of pain *Intensity*, confirming that participants rated the three intensity levels as progressively more painful. No modulation of *Task* was observed, neither as a main effect, nor in interaction with the *Intensity*. We also checked whether participants’ sensitivity to one’s pain changed as function of the performance of the preceding 3-Back condition, ad described by both IES and ITI values. Results are displayed in Table 2 and Figure 2B reports the results, confirming the main effect of stimulation *Intensity*, but also describing significant *IES*Intensity* and *ITI*Intensity* interactions. In particular, blocks associated with poorer performance led to significant decreased sensitivity to MP (*vs*. LP) stimulations, for both IES and ITI (Table 2 & Figure 2).

### Questionnaire Scores

We repeated the analyses of pain ratings by including the scores of relevant questionnaires of interest as covariates, in order to identify potential determinants of inter-individual differences in our effects. Specifically, we considered the Pain Catastrophizing Scale^73^, which is expected to tap key components involving attention and cognitive control of pain. As this questionnaire contains three subscales of interest (Rumination, Helplessness and Magnification), each of which could lead to plausible relevant effects, we report effects if associated with an α ≤ 0.0166 (corresponding to Bonferroni-corrected α ≤ for three multiple tests). We found no effects associated in any of the three sub-scores in any of the measures of Self- and Other-Pain.

We also tested whether scores of the Interpersonal Reactivity Index^23^, a standardized questionnaire of individual empathic traits, could influence the Other-Pain effects. As this questionnaire contains four subscales of interest (Personal Distress, Perspective Taking, Empathic Concern, Fantasy), we report effects if associated with an α ≤ 0.0125 (Bonferroni-corrected for four tests) for the Other-Pain condition. We found only a significant main effect of *Personal Distress* in the analysis of Pain intensity ratings, suggesting that individuals with higher scores are more prone to rate others’ expressions as more painful (*t*_(29.04)_ = 3.20, *p* = 0.003; *d* = 0.59; see Figure 3). No other main/interaction effect was associated with any of the questionnaire scores.

**Figure 3.**
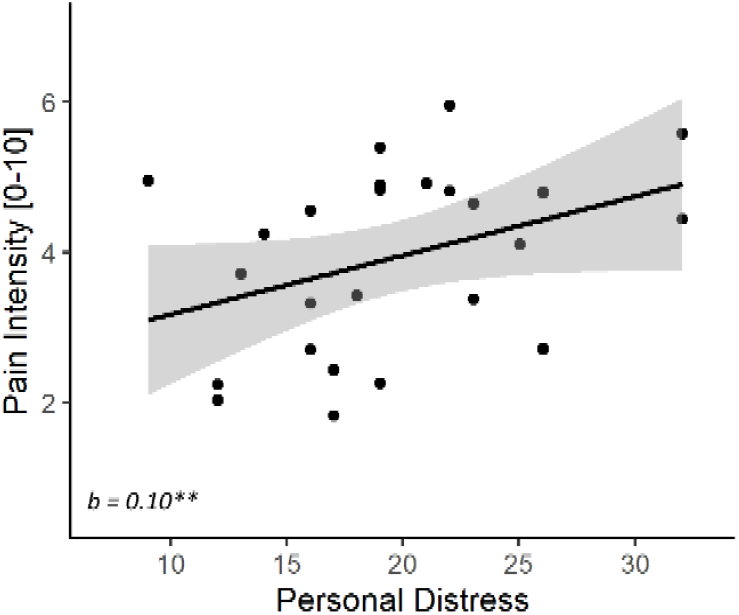
Experiment 1. Personal Distress effect on Other-Pain data. Individuals with higher scores on the Personal Distress subscale of IRI were more prone to rate others’ expressions as more painful. The relationship is described in terms of a linear regression line, associated with 95% confidence intervals area and data-points.

### Additional Measures

Supplementary Results report also details about the effects associated with physiological responses, and confidence ratings. Briefly, in the Self-pain condition all physiological measures (skin conductance response and heart rate variability) were significantly associated with a main effect of stimulation *Intensity*, reflecting a progressing increase of response, the stronger energy levels. We found neither an effect of *Task*, nor a modulation of participants’ performance (IES, ITI) on pain response. Instead, no significant effect was associated with the Other-pain condition. Finally, the analysis of confidence ratings revealed no significant effects.

## Experiment 2

Experiment 2 was carried out as a conceptual replication of Experiment 1, aimed at verifying whether the effects observed could be found also when manipulating cognitive exhaustion through another paradigm. For this reason, we run a new study which was identical to Experiment 1, except that the N-Back task was replaced with Stroop similar to that used in previous studies investigating effects of cognitive exhaustion on self-pain^65,68^.

### Methods

Unless stated otherwise, recruitment criteria, experimental set-up, procedure and data analyses were identical to those used in Experiment 1.

#### Participants

We recruited a total of 39 participants, six of which were subsequently excluded due to falling asleep during data acquisition (one participant) or lack of sensitivity to the nociceptive stimulation (five participants). We further excluded 8 participants who were not susceptible to the task manipulation. This led to a final sample of 25 individuals (14 females; aged 18 to 36, Mean =23.52, SD = 4.28 years). The exclusion/inclusion criteria were the same as in Experiment 1. In addition, as Experiment 2 involved verbal material, we recruited only French native speakers. The sample size was selected a priori to be identical to that from Experiment 1.

#### Stroop task

In Experiment 2, N-Back blocks were replaced by Stroop sessions, similar to those previously found having an after-effect on self-pain^65,68^. Each participant performed neutral and interference conditions of the numerical Stroop task^13^. Participants were informed that they would see sets of one to four identical words presented vertically on the screen. They were asked to count how many times a word was presented on the screen using buttons of the response box as quickly and accurately as possible, with a strong emphasis placed on not sacrificing accuracy for speed. Moreover, participants were asked not to use strategies to make the task easier (like blur words on the screen). In the interference condition, words stimuli were number words ‘one’, ‘two’, ‘three’ and ‘four’ – presented in French (‘un’, ‘deux’, ‘trois’, ‘quatre’) and they were always incongruent stimuli (word meaning and number of repetitions were different). In the neutral condition, words stimuli were neutral words matched in length with the number words: ‘an’, ‘drap’, ‘table’ and ‘quille’, corresponding in English to ‘year’, ‘sheet’, ‘table’ and ‘skittle’. All words were presented in capital letters. Trials started with a fixation cross (1000 ms) followed by a display with words to count which stayed on the screen until participants gave their response but no more than 1250 ms.

Prior to the main experiment, participants carried out a brief training session, in which they were administered 18 practice trials with neutral condition. During the training they received correctness feedback and were also informed to answer more quickly if they did not provide an answer within the allotted time. Feedback appeared for 2500 ms. In the main experiment, no feedback and speed-instruction were present. Furthermore, to control for order effects half of the participants (n = 13, 6 women) were administered first four blocks with neutral condition (2 “Self” and 2 “Other” in pseudorandomized order), followed by a break of about 5 minutes and then four blocks with interference condition (2 “Self” and 2 “Other” in pseudorandomized order). Half of the participants instead had the interference condition first (n = 12, 9 women).

## Results

### Task Demands

We first assessed whether participants exhibited higher difficulty in the demanding conditions. We focused on all participants who displayed adequate sensitivity to pain (N = 33; see Methods) and found that, on average, they displayed significant differences across the two conditions of interest in reaction times of correct responses (*t*_(32)_ = 5.53, *p* < 0.001; *d* = 0.98), and combined inverse efficiency score (IES, *t*_(32)_ = 4.41, *p* < 0.001; *d* = 0.78). However, although the tasks appeared to influence the performance of the overall population, few participants appeared non-susceptible to the manipulation. Figure 1 (right subplot) displays the individual inverse efficiency scores from the population, revealing that 8 participants showed comparable if not more proficient performance in the difficult condition. As in Experiment 1, the data from these 8 participants were excluded from subsequent analysis, thus leading to a final sample of N = 25.

### Pain Ratings

We then assessed whether sensitivity to nociceptive stimulation on one’s own body was influenced by the preceding task. Table 3 and Figure 4A reports results associated with pain intensity ratings, and revealed a significant main effect of stimulation *Intensity*, confirming that participants rated the three intensity levels as progressively more painful. We also found that participants’ ratings were influenced by the preceding task, in the shape of a *Task*Intensity* interaction. We further explored this interaction through simpler models testing effects of *Task* for each stimulation intensity level. The interaction was explainable by a significant decreased response to HP following Stroop interference *vs*. control (*t* = -2.79, *p* ≤ 0.006). In addition, there was a marginal increase in pain ratings for LP (*t*_(22.56)_ = 1.80, *p* = 0.085; *d* = 0.38), in line with what already observed in previous studies^65,68^. We also checked whether participants’ sensitivity to one’s pain changed as function of the performance of the preceding Stroop condition, by modelling participants’ IES from the previous block as a between-subjects continuous predictor. The results are displayed in Table 4, confirming the main effect of stimulation *Intensity*, but providing no evidence of an effect of IES, neither as main effect or in interaction with other variables.

**Table 3.**
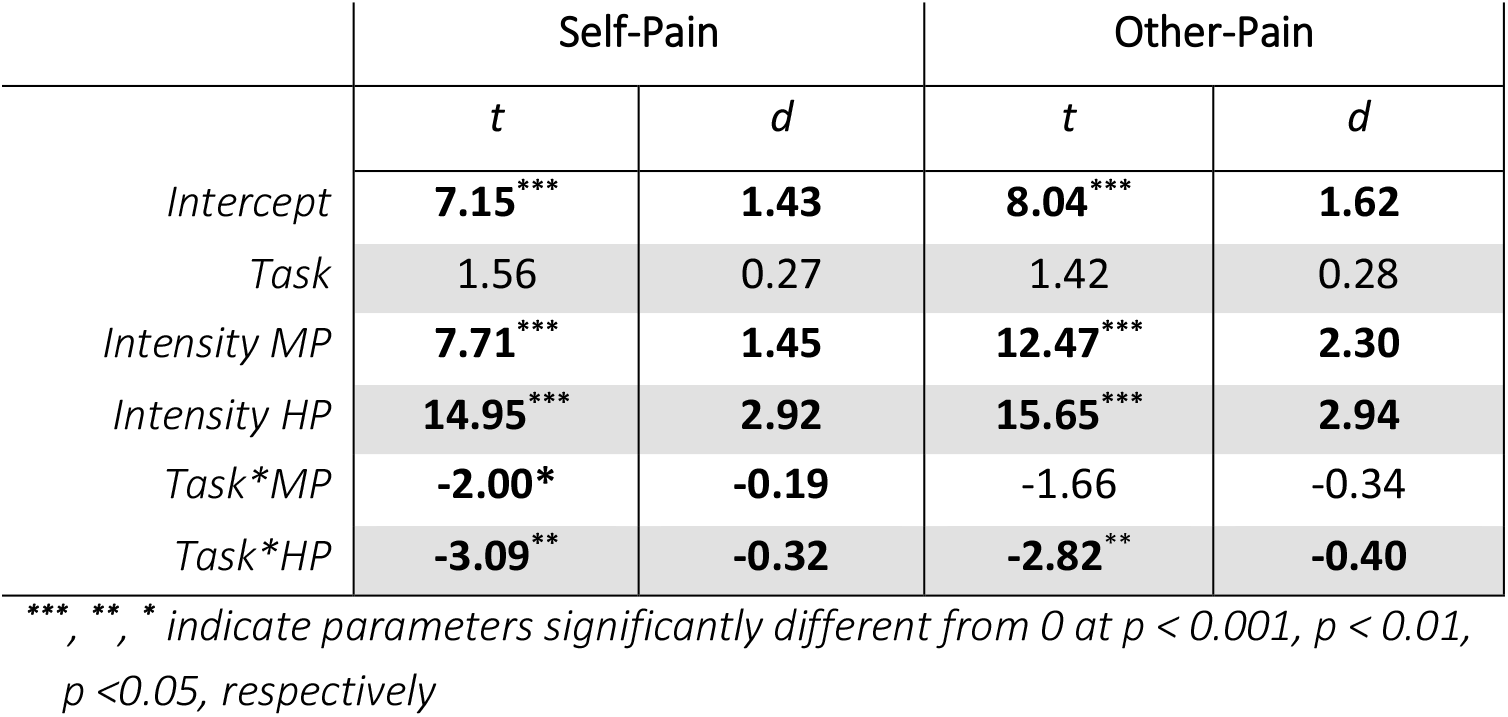
Experiment 2. Analysis of Task-effects on pain. We report the t-values and effect sizes (d) associated with parameter estimates from linear mixed model analyses testing effects of task on Self and Others’ pain. Significant effects are highlighted based on the corresponding p-value. MP = Medium Pain; HP = High Pain.

|  | Self-Pain |  | Other-Pain |  |
| --- | --- | --- | --- | --- |
|  | *t* | *d* | *t* | *d* |
| *Intercept* | **7.15\*\*\*** | **1.43** | **8.04\*\*\*** | **1.62** |
| *Task* | 1.56 | 0.27 | 1.42 | 0.28 |
| *Intensity MP* | **7.71\*\*\*** | **1.45** | **12.47\*\*\*** | **2.30** |
| *Intensity HP* | **14.95\*\*\*** | **2.92** | **15.65\*\*\*** | **2.94** |
| *Task\*MP* | **-2.00\*** | **-0.19** | -1.66 | -0.34 |
| *Task\*HP* | **-3.09\*\*** | **-0.32** | **-2.82\*\*** | **-0.40** |

\*\*\*, \*\*, \* indicate parameters significantly different from 0 at $p < 0.001$ , $p < 0.01$ , $p < 0.05$ , respectively

**Table 4.** Experiment 2. Analysis of Task Performance. We report the t-values and effect sizes (d) associated with parameter estimates from linear mixed model analyses testing effects of task performance and difficulty on Self and Others’ pain. Significant effects are highlighted based on the corresponding p-value. MP = Medium Pain; HP = High Pain; IES = Inverse Efficiency Score.

|  | Self-Pain: *IES* |  | Other-Pain: *IES* |  |
| --- | --- | --- | --- | --- |
|  | *t* | *d* | *t* | *d* |
| *Intercept* | **9.16<sup>\*\*\*</sup>** | **1.87** | **8.91<sup>\*\*\*</sup>** | **1.83** |
| *Intensity MP* | **5.58<sup>\*\*\*</sup>** | **1.06** | **8.11<sup>\*\*\*</sup>** | **1.63** |
| *Intensity HP* | **12.01<sup>\*\*\*</sup>** | **2.51** | **9.83<sup>\*\*\*</sup>** | **2.02** |
| *IES* | 0.90 | 0.10 | -1.20 | -0.14 |
| *IES\*MP* | -0.74 | -0.07 | -0.51 | -0.06 |
| *IES\*HP* | 0.09 | 0.01 | -0.15 | -0.02 |

<sup>\*\*\*</sup>, <sup>\*\*</sup>, <sup>\*</sup> indicate parameters significantly different from 0 at $p < 0.001$ , $p < 0.01$ , $p < 0.05$ , respectively

**Figure 4.**
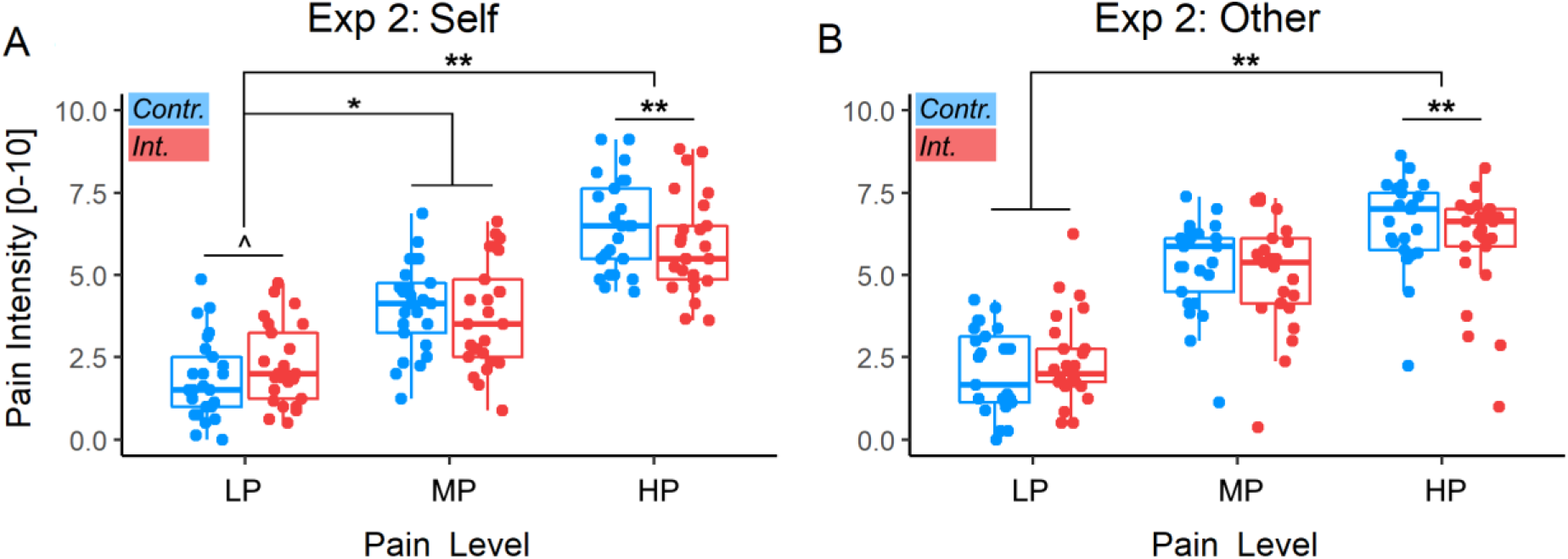
Experiment 2. Analysis of Task-effects on pain. Pain intensity Stimulation for Self-Pain (A) and Other-Pain data (B). Red boxplots and data refer to nociceptive stimulations occurring after the Interference [Int.] Stroop condition, whereas blue boxplots/data refer to stimulations following the easy Control [Contr.]. Box plots are described in terms of median (horizontal middle line), interquartile range (box edges), and overall range of non-outlier data (whiskers). Dots refer to individual average values associated to each condition, and are considered outliers if exceeding 1.5 inter-quartile ranges from the median. **, *, ^ refer to significant task main effects for a given pain stimulation level, or to significant interactions between Task and pain intensity at p < 0.01, p<0.05, and 0.05<p<0.1 respectively. LP = Low Pain; MP = Medium Pain; HP = High Pain.

Next, we analysed pain ratings associated with videos of patients displaying painful/painless facial expressions. Results are reported in Table 3 and Figure 4B and reveal a significant main effect of pain *Intensity*. As for the case of the Self-Pain ratings, we observed also *Task*Intensity* interaction. We further explored this interaction through simpler models testing effects of *Task* in each pain level, which was explained by a significant decreased response to HP following Stroop interference *vs*. control (*t*_(67.11)_ = - 2.97, *p* = 0.004; *d* = -0.36; Figure 2B). Finally, we also checked whether participants’ sensitivity to other’s expressions changed as function of the performance of the preceding depleting task, but found no effect associated with IES.

### Additional Measures

As for Experiment 1, we repeated the analyses of pain ratings by including the scores of the Pain Catastrophizing Scale^73^, and Interpersonal Reactivity Index^23^, as predictor. No significant effect was found. Supplementary Results report also details about the effects associated with physiological responses, and confidence ratings. Briefly, in the Self-pain condition all physiological measures (skin conductance response and heart rate variability) were significantly associated with a main effect of stimulation *Intensity*. Furthermore, we found that physiological measures were also modulated by the preceding task, in the shape of a *Task*Intensity* interaction (for skin conductance level) or a *IES*Intensity* interaction (for hearth rate variability). In both cases, the interaction is explainable in terms of decreased response to HP following a more challenging block. No significant effect in physiological responses was associated with the Other-pain condition. Finally, the analysis of confidence ratings revealed no significant effects.

## Discussion

We tested whether cognitive exertion influences the evaluation of subsequent one’s and other’s pain. For this purpose, we employed two cognitively demanding tasks, a working memory (*Exp. 1*: N-Back)^12^ and a cognitive control task (*Exp. 2*: Stroop)^26,62,65^, which both decreased the sensitivity (measured through intensity ratings) to pain on one’s own body, especially for medium and high stimulations. Consistent effects were also observed for galvanic skin response and heart rate variability (see supplements). These effects were mirrored by a comparable decrease in subsequent assessment of facial expressions, with participants underrating those video-clips displaying the middle (*Exp. 1*: N-Back) or highest pain (*Exp. 2*: Stroop). This was observed either by comparing the effects of demanding condition to those associated with an easier control condition (*Exp. 2*: Stroop), or by modelling linearly the difficulty/performance of each demanding block (*Exp. 1*: N-Back). Finally, and specifically for the Stroop, we also found that cognitive control led to a mild hypersensitization towards self-directed low-pain stimulations (as previously suggested)^65,68^. In both experiments, and in both self- and other-assessment, task modulations on pain sensitivity ranged between a Cohen’s *d* of 0.17 to 0.40, reflecting a magnitude within the small to medium range^19^. Overall, our results show that cognitive exertion alters the assessment others’ suffering, partly mirroring to what happens for first-hand pain experiences.

### Stroop effects on Self-pain

The Stroop task influenced self-pain experience, by increasing marginally sensitivity for LP stimuli, and decreasing it for HP ones. Although, to our knowledge, Stroop aftereffects on pain have never been tested as function of stimulation intensity, our findings may explain the heterogeneity in the literature, as previous research reported hyperalgesic effects of the Stroop for medium-low noxious stimulations (ratings corresponding to ∼4 on our scale)^65,68^, whereas others described hypoalgesia for more intense events (∼6)^33^. Traditionally, hyperalgesia following demanding tasks is interpreted in light of impaired top-down regulatory processes, which are elicited following nociceptive events to downplay aversive experiences, and promote coping reactions^65,68^. This interpretation fits well our findings with mild stimulations, but it does not account for what observed for intense nociceptive stimulations.

Our results reveal that cognitive exertion has a differential impact on mild and high pain stimulations, for which we suggest two possible interpretations. First, regulatory processes may be engaged in an active control of moderately (non-life-threatening) painful situations to allow the pursuit of one’s everyday activity. Hence, for moderate levels of pain, increased attentional demands would impair efficient pain downregulation. Instead, when experiencing intense (or enduring) pain, the body may enter in a shock-like state in which suffering responses are automatically dulled^6,36^. Accordingly, the more a noxious stimulus is intense, the more pain regulation might shift from deliberate to automatic, thus leading to opposing depletion effects for high *vs*. low intensity. A second plausible interpretation would imply that cognitive exertion does not influence uniquely top-down regulatory processes, but interferes directly with the representation of pain. Indeed, recent accounts suggest that pain experience could be best interpreted within a Bayesian framework, where the brain estimates the (posterior) probability of potential body damage, based on the integration of sensory inputs and prior representations^10,51,52,61,62,64,74,75,84^. Within this framework, pain should be considered the result of an active inferential process which, under limited resources, could prove challenging for individuals, and lead poorer discrimination between low and high stimulations. This last interpretation would effectively fit all our findings.

In either case, our data provide first clear evidence that cognitive control aftereffects on pain change as function of the intensity of the noxious stimulation, thus helping disambiguate an otherwise mixed literature.

### N-Back effects on Self-pain

As for the effects of a memory task on self-pain, we found that participants’ performance in each 3-Back block was linearly coupled with the subsequent rating of medium/high pain stimulations: specifically, the poorer the performance (high IES) the higher the subsequent hypoalgesia (Figure 3A). Such effect could be interpreted as a result of cognitive depletion (as for the case of the Stroop), but only under the assumption that poor performance underlies a high level of cognitive fatigue already present during the task, and extending on the subsequent nociceptive stimulations.

An alternative interpretation spawns from motivational models of depletion and cognitive fatigue, according to which task subjective difficulty interacts with individual commitment in mobilizing effort^66,67,85^. Within this framework, a task perceived as too difficult would lead individuals to disengage, thus impacting only minimally the cognitive resources. Hence, participants with poor performance (high IES) should not necessarily have a high cognitive strain, but might have not committed to the task and, paradoxically, would have more resources available during the subsequent painful stimulation. We believe that this interpretation does not fit entirely our results. Indeed, while it is true that participants who disengaged from the task should be characterized by low proficiency (high IES), they should also be associated with low difficulty (short ITI). Indeed, as the task difficulty was continuously adapted to stabilize as much as possible performance across the population (see Methods), individuals who “gave up” should have rapidly converged towards the easiest task parameters. This was not observed in our study where, in the Self-Pain condition, the ITI did not influence participants’ sensitivity to pain. For this reason, participants’ high IES in our study most likely reflects proficiency fluctuations while they are actively engaged in the task.

Overall, although the results from the N-Back task are not entirely comparable with those from the Stroop, both sets of results converge in showing that task aftereffects impact the sensitivity to pain differentially as function of the noxious stimulation, with decreased sensitivity towards medium/high events.

### Effects on Other-pain

The present study allowed us to compare opposing predictions between two accounts: the *Broaden-and-Build theory* indirectly suggested that cognitive exertion should influence self and other’s pain differentially, by increasing sensitivity for the former, at the expense of the latter; instead, *Embodied* accounts positing shared representation between self and others’ pain, predict that cognitive exertion might affect judgments of others’ pain analogously to self-pain. Our results provide support to some predictions from both accounts. In line with the *Broaden-and-Build* theory, restrictions of one’s cognitive resources did decrease the sensitivity of medium/high painful expressions in both N-Back (as function of task performance and difficulty) and Stroop (relative to a neutral control condition). However, and most importantly, all the task-aftereffects observed in for other’s pain were observed also for the self-pain, with astonishing similarity in terms of the way in which the previous task was modelled (Task factor in Stroop, IES regressor in N-Back) and on the pain levels implicated (HP for Stroop, MP for N-Back). Hence, whatever the effects of each cognitive task on self-pain, similar effects were present also for the observation of others’ painful expression, thus providing strong support to the Embodied account.

Regardless of the winning model, our study sheds light on the role of mental fatigue in pain diagnosis, with relevant translational implications for healthcare practice. Physicians and nurses are often tasked with assessing patients’ pain under tiring working schedules. As such, they often underestimate patients’ pain^57,58^, a tendency present even during early university training^20,26^. As our data suggest some convergence between N-Back and Stroop in their negative effects on pain assessment, we are incline to assume that our effects might be generalizable also to other kinds of cognitive load. However, it is unclear whether these include also those condition most relevant in clinical practice, such as sleep loss, extreme working schedules, and/or being engaged in multiple tasks at the same time. Future studies will need to extend our findings to other forms of cognitive exhaustion of more ecological values in clinical practice (e.g., through sleep deprivation paradigms).

### Limitations of the study and conclusive remarks

In our study, we excluded participants based on low pain sensitivity or non-susceptibility to the Stroop/N-Back. Although this criterion was established *a priori*, and data collection was conducted until a desired number of suitable participants was obtained, two related limitations need to be underscored. The first is the sub-optimal nature of the task manipulation and pain intensity calibration, which might be vulnerable to factors like training, habituation, desensitization, etc. The second is the final sample, which might not be representative of the overall population, but only of those individuals most sensitive to pain and less proficient in the tasks employed. Hence, we might have excluded those individuals with high cognitive resources and regulatory abilities, who were able to perform difficult tasks effortlessly and/or might be extremely efficient at tempering down the aversive effects of painful events in their body.

Furthermore, differently from the case of the Stroop, N-Back aftereffects on pain sensitivity were found only when modelling individual IES as continuous predictor, but not in relation to the *a priori* designed control 0-Back. We suggest caution interpreting differences between the two experiments, as the two tasks were selected from previous research due to their established efficacy in modulating self-pain experience^12,68^, but they have never been matched with one another in terms of difficulty. However, one possible explanation for this observation could be that memory tasks might be less suited than Stroop for inducing a depletion effect, as suggested by a recent meta-analysis^14^. Cognitive control might be required also in the 0-Back control condition, where participants are asked to only respond to the presentation of a specific target letter (similarly to a “go/no-go” paradigm^80^), thus potentially questioning the effectiveness of this condition as a control for cognitive control effects. Future studies will need to use a more suitable control (e.g. 1-Back) to test whether any discrepancy between working memory and inhibitory control in their relation to pain might hold. Finally, another point of divergence between the two studies was the susceptibility of other pain ratings to individual’s personal distress trait from the IRI scale. This result would be consistent with previous research who linked this trait with the assessment of other people’s pain^27^, although it is unclear why it was observed only in Experiment 1, and not in Experiment 2 where the same videos were preceded by the Stroop task.

Notwithstanding these limitations, our study demonstrates that the aftereffect of cognitive exertion on others’ pain judgment decreases the sensitivity towards the most intense stimulations, similarly to what observed in first-hand experiences. Healthcare personnel should be aware that high workload and strong cognitive fatigue might alter their diagnostic abilities.

## Acknowledgments

We would like to thank Dr. Flavia Mancini and the team of Prof. Luis Garcia-Larrea (Dr. Maud Frot and Dr. Caroline Perchet) for all their assistance involving the nociceptive stimulations. We also thank Gwénaël Birot, Roberto Martuzzi, Leyla Loued-Khenissi and Loan Mattera for their support in data collection, Ben Meuleman and Lia Antico for their assistance in data analysis, and the Campus Biotech for the use of equipment and space.

