## Supplementary material for "Cognitive exertion affects the appraisal of one’s own and other people’s pain"

### Supplementary Information

#### Supplementary Methods

##### Physiologic Responses

During the main experiment, we continuously recorded physiological responses to stimuli from their left hand, such as skin conductance (EDA) and heart rate. These physiological measures were acquired using the Biopac System and Acknowledge software. Electrodermal activity was recorded from two disposable GSR/EDA electrodes (EL507, Neurospec AG) placed on the middle and ring fingers, while cardiac pulse was recorded placing a transducer on the index finger (Biopac System).

###### *Skin conductance*

Electrodermal activity data were first subjected to a low-pass filter (cut-off frequency: 5Hz) to account for movement-related artefacts. The filtered signal was then processed and analysed with Ledalab[3], a free Matlab-based toolbox. More specifically, the time course was down sampled to 50 Hz, smoothed (adaptive Gaussian), and visually inspected for potential remaining movement artefacts, which were corrected through spline interpolation. The resulting signal was then de-convolved using continuous decomposition analysis, which separates traces into tonic (slowly changing skin conductance level) and phasic (rapidly changing response) activity components. For the purpose of this analysis, we considered as reliable skin conductance response (SCR), a cumulative increase of phasic activity occurring between 1 and 5 s from the painful event (either on one's body or through video) and exceeding 0.01 μS. These single trial estimates of SCR were square-root transformed to improve compliance with normal distribution and fed to the same statistical pipeline used for the analysis of the behavioural measures.

###### *Cardiac response*

As for pulse, cardiac response was band-pass filtered (between 10–30Hz), electrocardiographic R waves were detected offline, and intervals between heartbeats were used to estimate Heart Rate Variability (HRV) over a time-window of 14 seconds from the onset of the painful stimulation/video. More specifically, we calculated single trial estimates of the Root-Mean-Squared Sequential Difference (RMSSD)[64], which is a measure used to quantify the amount of HRV observed during monitoring periods that generally may range from <1 min to >24 h. HRV and its indices like the RMSSD have been shown to be reliable indexes of physiologic response to pain[35] and they have been proposed as measures of cognitive fatigue and self-regulatory strength[45,60]. This time-window was chosen as it captures modulations associated with painful events and ratings, without never exceeding the onset of the subsequent trial. RMSSD measures were then fed to the same analytical pipeline used for the other measures.

#### Supplementary Results

The analyses of confidence ratings and physiological measures was carried out under the same linear mixed models framework used for the pain intensity ratings. Please check Supplementary Figures and Tables for more details

#### Supplementary Tables

| | Exp 1: *Self-Pain* | | | | | |
|---|---|---|---|---|---|---|
| | *Conf. R.* | | *SCR* | | *HRV* | |
| | *t* | *d* | *t* | *d* | *t* | *d* |
| *Intercept* | **29.80**$^{***}$ | **5.93** | **6.23**$^{***}$ | **1.27** | **7.45**$^{***}$ | **1.53** |
| *Task* | 0.63 | 0.12 | 0.58 | 0.12 | 0.05 | 0.01 |
| *Intensity MP* | -1.29 | -0.21 | 0.38 | 0.07 | 0.21 | 0.03 |
| *Intensity HP* | -1.09 | -0.20 | **4.88**$^{***}$ | **0.99** | 0.83 | 0.15 |
| *Task*MP* | 0.09 | 0.01 | 1.16 | 0.21 | -0.29 | -0.06 |
| *Task*HP* | -0.25 | -0.03 | -0.68 | -0.13 | 0.50 | 0.10 |

$^{***}$, $^{**}$, $^{*}$ *indicate parameters significantly different from 0 at p < 0.001, p < 0.01, p <0.05, respectively*

***Table S1***. *Experiment 1. Analysis of Task-effects on self-pain. We report the t-values and effect sizes (d) associated with parameter estimates from linear mixed model analyses run on each experiment. Significant effects are highlighted based on the corresponding p-value (see legend). Conf. = confidence; R. = rating; SCR = skin conductance response; HRV = Heart Rate Variability; MP = Medium Pain; HP = High Pain*

| | *Exp 1*: Self-Pain, *IES* analysis | | | | | |
|---|---|---|---|---|---|---|
| | *Conf. R.* | | *SCR* | | *HRV* | |
| | *t* | *d* | *t* | *d* | *t* | *d* |
| *Intercept* | **32.52*****| **6.54** | **7.89***** | **1.64** | **7.54***** | **1.54** |
| *Intensity MP* | -1.37 | -0.09 | **2.25*** | **0.36** | -0.21 | -0.04 |
| *Intensity HP* | -1.81 | -0.12 | **5.65***** | **1.19** | 1.72 | 0.35 |
| *IES* | 1.01 | 0.07 | 1.72 | 0.13 | -0.70 | -0.06 |
| *IES*MP* | 0.43 | 0.03 | -1.09 | -0.11 | 0.09 | 0.01 |
| *IES*HP* | -1.41 | -0.09 | -1.24 | -0.14 | 0.41 | 0.04 |

****, **, * indicate parameters significantly different from 0 at p < 0.001, p < 0.01, p <0.05, respectively*

***Table S2***. *Experiment 1. Analysis of IES on Self-Pain. We report the t-values and effect sizes (d) associated with parameter estimates from linear mixed model analyses run on each experiment. Significant effects are highlighted based on the corresponding p-value. Conf. = confidence; R. = rating; MP = Medium Pain; HP = High Pain; IES = Inverse Efficiency Score; SCR = skin conductance response; HRV = Heart Rate Variability.*

| | *Exp 1*: Self-Pain, *ITI* analysis | | | | | |
|---|---|---|---|---|---|---|
| | *Conf. R.* | | *SCR* | | *HRV* | |
| | *t* | *d* | *t* | *d* | *t* | *d* |
| *Intercept* | **32.63***** | **6.57** | **8.14***** | **1.67** | **7.75***** | **1.60** |
| *Intensity MP* | -1.36 | -0.09 | **2.26*** | **0.31** | -0.21 | -0.04 |
| *Intensity HP* | -1.72 | -0.32 | **5.46***** | **1.12** | 1.72 | 0.35 |
| *ITI* | -1.05 | -0.07 | -1.45 | -0.10 | -1.57 | -0.11 |
| *ITI*MP* | 1.23 | 0.08 | 1.96 | 0.14 | 1.97 | 0.14 |
| *ITI*HP* | 0.79 | 0.06 | 0.89 | 0.07 | 0.50 | 0.04 |

****, **, * indicate parameters significantly different from 0 at p < 0.001, p < 0.01, p <0.05, respectively*

***Table S3***. *Experiment 1. Analysis of ITI on Self-Pain. We report the t-values and effect sizes (d) associated with parameter estimates from linear mixed model analyses run on each experiment. Significant effects are highlighted based on the corresponding p-value. Conf. = confidence; R. = rating; MP = Medium Pain; HP = High Pain; ITI = Inter-trial Interval; SCR = skin conductance response; HRV = Heart Rate Variability.*

| | Exp 1: *Other-Pain* | | | | | |
|---|---|---|---|---|---|---|
| | *Conf. R.* | | *SCR* | | *HRV* | |
| | *t* | *d* | *t* | *d* | *t* | *d* |
| *Intercept* | **13.43**$^{***}$ | **2.76** | **3.13**$^{***}$ | **0.99** | **7.62**$^{***}$ | **1.54** |
| *Task* | 0.42 | 0.08 | -1.62 | -0.27 | -0.13 | -0.02 |
| *Intensity MP* | -1.17 | -0.15 | -1.43 | -0.26 | -0.13 | -0.02 |
| *Intensity HP* | 0.15 | 0.02 | -0.59 | -0.22 | -0.12 | -0.02 |
| *Task*MP* | -0.48 | -0.07 | 1.54 | 0.10 | -0.28 | -0.03 |
| *Task*HP* | 0.55 | 0.09 | 0.54 | 0.04 | 0.14 | 0.01 |

$^{***}$, $^{**}$, $^{*}$ *indicate parameters significantly different from 0 at p < 0.001, p < 0.01, p <0.05, respectively*

***Table S4***. *Experiment 1. Analysis of Task-effects on other-pain. We report the t-values and effect sizes (d) associated with parameter estimates from linear mixed model analyses run on each experiment. Significant effects are highlighted based on the corresponding p-value. Conf. = confidence; R. = rating; MP = Medium Pain; HP = High Pain; SCR = skin conductance response; HRV = Heart Rate Variability.*

| | ***Exp 1*: Other-Pain, *IES* analysis** | | | | | |
|---|---|---|---|---|---|---|
| | *Conf. R.* | | *SCR* | | *HRV* | |
| | *t* | *d* | *t* | *d* | *t* | *d* |
| *Intercept* | **13.68**$^{***}$ | **2.83** | **5.40**$^{***}$ | **1.05** | **6.61**$^{***}$ | **1.38** |
| *Intensity MP* | -1.66 | -0.27 | -0.65 | -0.04 | -0.45 | -0.08 |
| *Intensity HP* | 0.77 | 0.16 | -1.24 | -0.11 | 0.06 | 0.01 |
| *IES* | -1.28 | -0.09 | -0.79 | -0.06 | -1.76 | -0.13 |
| *IES*MP* | -0.57 | -0.05 | 1.94 | 0.12 | 0.66 | 0.07 |
| *IES*HP* | 0.50 | 0.06 | 0.95 | 0.06 | 0.46 | 0.04 |

$^{***}$, $^{**}$, $^{*}$ *indicate parameters significantly different from 0 at p < 0.001, p < 0.01, p < 0.05, respectively*

***Table S5***. *Experiment 1. Analysis of IES on Other-Pain. We report the t-values and effect sizes (d) associated with parameter estimates from linear mixed model analyses run on each experiment. Significant effects are highlighted based on the corresponding p-value. Conf. = confidence; R. = rating; MP = Medium Pain; HP = High Pain; IES = Inverse Efficiency Score; SCR = skin conductance response; HRV = Heart Rate Variability.*

| | ***Exp 1*: Other-Pain, *ITI* analysis** | | | | | |
|---|---|---|---|---|---|---|
| | *Conf. R.* | | *SCR* | | *HRV* | |
| | *t* | *d* | *t* | *d* | *t* | *d* |
| *Intercept* | **14.28*****| **2.92** | **5.33***** | **1.04** | **6.86***** | **1.40** |
| *Intensity MP* | -1.63 | -0.25 | -0.63 | -0.12 | -0.45 | -0.08 |
| *Intensity HP* | 0.81 | 0.17 | -1.22 | -0.10 | 0.11 | 0.01 |
| *ITI* | -1.16 | -0.07 | -0.02 | -0.01 | -0.91 | -0.06 |
| *ITI*MP* | 0.46 | 0.03 | 0.47 | 0.04 | 0.35 | 0.02 |
| *ITI*HP* | 0.28 | 0.02 | -0.67 | -0.04 | -0.86 | -0.06 |

***, **, * *indicate parameters significantly different from 0 at p < 0.001, p < 0.01, p < 0.05, respectively*

***Table S6****. Experiment 1. Analysis of ITI on Other-Pain. We report the t-values and effect sizes (d) associated with parameter estimates from linear mixed model analyses run on each experiment. Significant effects are highlighted based on the corresponding p-value. Conf. = confidence; R. = rating; MP = Medium Pain; HP = High Pain; ITI = Inter-trial Interval; SCR = skin conductance response; HRV = Heart Rate Variability.*

| | Exp 2*: Self-Pain* | | | | | |
|---|---|---|---|---|---|---|
| | *Conf. R.* | | *SCR* | | *HRV* | |
| | *t* | *d* | *t* | *d* | *t* | *d* |
| *Intercept* | **30.79***** | **6.25** | **6.46***** | **1.36** | **8.47***** | **1.55** |
| *Task* | 0.31 | 0.06 | 0.72 | 0.14 | 0.78 | 0.09 |
| *Intensity MP* | -1.57 | -0.32 | **3.81***** | **0.63** | -0.24 | -0.02 |
| *Intensity HP* | -0.38 | -0.08 | **6.77***** | **1.45** | **2.19*** | **0.25** |
| *Task*MP* | -0.17 | -0.03 | -1.18 | -0.11 | 0.88 | 0.04 |
| *Task*HP* | -0.38 | -0.08 | **-2.22*** | **-0.33** | -0.81 | -0.04 |

***, **, * *indicate parameters significantly different from 0 at p < 0.001, p < 0.01, p <0.05, respectively*

***Table S7****. Experiment 2. Analysis of Task-effects on self-pain. We report the t-values and effect sizes (d) associated with parameter estimates from linear mixed model analyses run on each experiment. Significant effects are highlighted based on the corresponding p-value (see legend). Conf. = confidence; R. = rating; SCR = skin conductance response; HRV = Heart Rate Variability; MP = Medium Pain; HP = High Pain*

| | ***Exp 2*: Self-Pain, *IES* analysis** | | | | | |
|---|---|---|---|---|---|---|
| | *Conf. R.* | | *SCR* | | *HRV* | |
| | *t* | *d* | *t* | *d* | *t* | *d* |
| *Intercept* | **31.99**$^{***}$ | **6.49** | **5.23**$^{***}$ | **1.07** | **6.89**$^{***}$ | **1.40** |
| *Intensity MP* | -1.29 | -0.26 | 1.41 | 0.18 | 0.85 | 0.18 |
| *Intensity HP* | -0.70 | -0.15 | **5.31**$^{***}$ | **1.07** | 1.05 | 0.21 |
| *IES* | 0.89 | 0.07 | 1.89 | 0.15 | 1.40 | 0.11 |
| *IES*MP* | -1.78 | -0.16 | -0.33 | -0.03 | -0.59 | -0.08 |
| *IES*HP* | -0.97 | -0.08 | 0.44 | 0.06 | **-2.28**$^{*}$ | **-0.28** |

$^{***}$, $^{**}$, $^{*}$ indicate parameters significantly different from 0 at $p < 0.001$, $p < 0.01$, $p <0.05$, respectively

***Table S8**. Experiment 2. Analysis of IES on Self-Pain. We report the t-values and effect sizes (d) associated with parameter estimates from linear mixed model analyses run on each experiment. Significant effects are highlighted based on the corresponding p-value. Conf. = confidence; R. = rating; MP = Medium Pain; HP = High Pain; IES = Inverse Efficiency Score; SCR = skin conductance response; HRV = Heart Rate Variability.*

| | Exp 2*: Other-Pain* | | | | | |
|---|---|---|---|---|---|---|
| | *Conf. R.* | | *SCR* | | *HRV* | |
| | *t* | *d* | *t* | *d* | *t* | *d* |
| *Intercept* | **15.14**$^{***}$ | **3.09** | **4.36**$^{***}$ | **0.88** | **7.90**$^{***}$ | **1.58** |
| *Task* | 0.10 | 0.02 | -0.54 | -0.05 | 0.70 | 0.14 |
| *Intensity MP* | 0.02 | 0.01 | 0.10 | 0.01 | 0.08 | 0.01 |
| *Intensity HP* | **2.62**$^{*}$ | 0.50 | 1.29 | 0.23 | 0.26 | 0.01 |
| *Task*MP* | 0.32 | 0.06 | 0.86 | 0.15 | 0.14 | 0.03 |
| *Task*HP* | -0.88 | -0.16 | 0.35 | 0.05 | 0.32 | 0.03 |

$^{***}$, $^{**}$, $^{*}$ indicate parameters significantly different from 0 at $p < 0.001$, $p < 0.01$, $p <0.05$, respectively

***Table S9**. Experiment 2. Analysis of Task-effects on Other-pain. We report the t-values and effect sizes (d) associated with parameter estimates from linear mixed model analyses run on each experiment. Significant effects are highlighted based on the corresponding p-value (see legend). Conf. = confidence; R. = rating; SCR = skin conductance response; HRV = Heart Rate Variability; MP = Medium Pain; HP = High Pain*

| | Exp 2: Stroop performance (*IES*) | | | | | |
|---|---|---|---|---|---|---|
| | *Conf. R.* | | *SCR* | | *HRV* | |
| | *t* | *d* | *t* | *d* | *t* | *d* |
| *Intercept* | **14.66***** | **3.00** | **3.90***** | **0.79** | **8.18***** | **1.67** |
| *Intensity MP* | 0.57 | 0.07 | 1.33 | 0.25 | 0.26 | 0.05 |
| *Intensity HP* | 1.63 | 0.33 | **2.10*** | **0.28** | 0.64 | 0.06 |
| *IES* | 0.16 | 0.01 | -1.04 | -0.08 | 1.17 | 0.12 |
| *IES*MP* | 0.08 | 0.01 | 0.91 | 0.12 | -1.64 | -0.23 |
| *IES*HP* | -1.09 | -0.16 | 0.61 | 0.06 | 0.24 | 0.02 |

****, **, * indicate parameters significantly different from 0 at p < 0.001, p < 0.01, p < 0.05, respectively*

***Table S10**. Experiment 2. Analysis of IES on Other-Pain. We report the t-values and effect sizes (d) associated with parameter estimates from linear mixed model analyses run on each experiment. Significant effects are highlighted based on the corresponding p-value. Conf. = confidence; R. = rating; MP = Medium Pain; HP = High Pain; IES = Inverse Efficiency Score; SCR = skin conductance response; HRV = Heart Rate Variability.*

#### Supplementary Figure

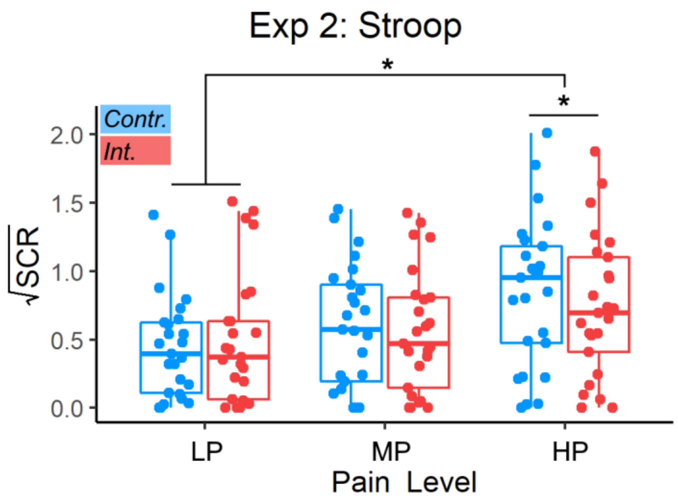


***Figure S1***. *Experiment 2. Analysis of Task-effects on skin conductance response (SCR) response to Self-Pain. Red boxplots and data refer to nociceptive stimulations occurring after the Interference [Int.] Stroop condition, whereas blue boxplots/data refer to stimulations following the easy Control [Contr.]. Box plots are described in terms of median (horizontal middle line), interquartile range (box edges), and overall range of non-outlier data (whiskers). Dots refer to individual average values associated to each condition, and are considered outliers if exceeding 1.5 inter-quartile ranges from the median. * refers to significant task main effects for a given pain stimulation level, or to significant interactions between Task and pain intensity at p<0.05. LP = Low Pain; MP = Medium Pain; HP = High Pain, Contr. = Stroop Control condition; Int. = Stroop Interference condition. SCR: Skin Conductance Response.*

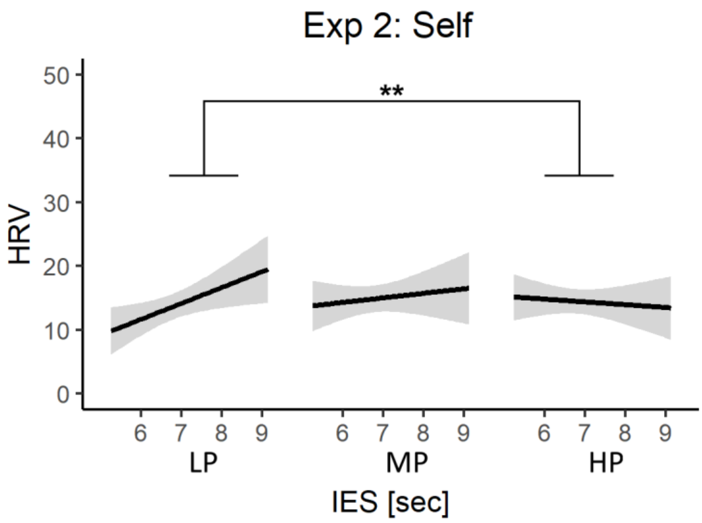


***Figure S2***. *Experiment 1. Analysis of Task Performance on cardiac responses of Self-Pain. For each pain level, the relationship between IES and pain ratings is described through a linear regression line with 95% confidence interval area. "**", to significant interactions between IES and pain intensity at p < 0.01. IES = Inverse Efficiency Score; LP = Low Pain; MP = Medium Pain; HP = High Pain; HRV = Heart Rate Variability.*
